# Reproductive microbial diversity is associated with competitive phenotypes in socially polyandrous jacanas

**DOI:** 10.1101/2025.02.05.636638

**Authors:** Jennifer L. Houtz, Kimberly A. Acosta, Mae Berlow, Sara E. Lipshutz

**Author notes:** Corresponding author: 130 Science Drive, Durham, NC, 27708.

## Abstract

The composition of host-associated microbial communities may reflect the overall status of the host, including physiology and reproductive success. New hypotheses suggest that reproductive microbiomes can mediate sexually selected phenotypes, which may be particularly important for species with mating systems that feature strong sexual selection. These dynamics have been particularly understudied in female animals. Using 16S rRNA sequencing, we compared the cloacal microbiome of females and males from two socially polyandrous bird species that vary in the strength of sexual selection, *Jacana spinosa* (Northern Jacana) and *J. jacana* (Wattled Jacana). We hypothesized that the strength of sexual selection would shape cloacal microbial diversity, such that the more polyandrous *J. spinosa* would have a more diverse microbiome, and that microbiomes would be more diverse in females than in males. If the reproductive microbiome is indicative of competitive status, we also hypothesized that cloacal microbial diversity would be associated with competitive traits, including plasma testosterone levels, body mass, or weaponry. We found a positive relationship between microbial alpha diversity and testosterone and weaponry in female *J. spinosa*. We found no differences in microbial alpha diversity between species or sexes, but we did find that microbial beta diversity significantly differed between species. Our results suggest that the cloacal microbiome may be a key component of the competitive phenotype in socially polyandrous jacanas, supporting the hypothesis that the interplay between sexual selection and microbial communities can shape the physiology and fitness of wild avian hosts.

**LAY SUMMARY:**

- The reproductive microbiome may be a key component of the competitive phenotype
- We examined cloacal microbial diversity in females and males of two *Jacana* species
- Microbial alpha diversity is associated with testosterone and weaponry in female *J. spinosa*
- Microbial beta diversity differed between *J. spinosa* and *J. jacana*

## INTRODUCTION

Almost all multicellular organisms host microbial communities, which consist of microorganisms (i.e., bacteria, viruses, unicellular algae, protozoans, and fungi) and their metagenomes - known as the ‘microbiome’ (Marchesi & Ravel, 2015). The composition of host microbiomes varies with diverse internal and external factors, including host genetics (Goodrich et al., 2014), diet (Bodawatta et al., 2022), and social interactions (Tung et al., 2015). The microbiome is not only shaped by its host; it also influences processes within the host, including digestion, metabolism, and protection from infection (Grond et al., 2018). Following the theory on biodiversity-ecosystem function, greater microbial diversity may increase its functionality in the host. For instance, hosts with higher microbial diversity may have a broader uptake of nutrients (Tennant et al., 1971), lower risk of immunodeficiency (Le Chatelier et al., 2013) and disease (Minamoto et al., 2015), and more resistance to invasion by microbial pathogens (Spragge et al., 2023). Because of the myriad ways microbiomes can impact host health, microbial diversity and composition are potentially an essential component of organismal fitness (but see (Williams et al., 2024)).

A recent focus in microbial ecology and evolution is the reproductive microbiome, which consists of microorganisms within structures, organs, fluids, or tissues of the host reproductive tract (Rowe et al., 2020). Though past work has emphasized sexually transmitted pathogens (Lombardo, 1998; Sheldon, 1993), the reproductive microbiome can affect a variety of reproductive traits including the host’s fertility (Williams et al., 2019), reproductive compatibility (Grieves et al., 2019), and hatching success (Van Dongen et al., 2019). Patterns of sexual transmission may influence the composition of the reproductive microbiome, as mating can facilitate the transfer of microbiota from one partner to another (Kulkarni & Heeb, 2007; Rowe et al., 2020). One consequence of this microbial transmission is that mating partners may converge to have similar reproductive microbiomes (Pruter et al., 2023; White et al., 2010). Another consequence is that females with multiple sexual partners may have higher reproductive microbiome diversity than monogamous females (MacManes, 2011; White et al., 2011). Thus, the reproductive microbiome may be impacted by the magnitude and direction of sexual selection, including variation in mating behavior from monogamy to polygamy (Rowe et al., 2020).

Reproductive microbiomes may also serve as honest signals of individual quality that indicate internal physiological states. Wild nonmodel organisms, including birds, have been valuable for examining the relationships between microbial diversity and fitness-related traits (Grond et al., 2018; Hird, 2017). For instance, a study of northern cardinals (*Cardinalis cardinalis*) found that ornamentation and body condition were positively correlated with cloacal microbiome diversity (Slevin et al., 2024). In rufous collared sparrows (*Zonotrichia capensis*), cloacal microbiome diversity was positively correlated with testosterone levels (Escallón et al., 2016). In birds and most other species, studies have focused on the relationship between the microbiome and sexually selected traits in males, leaving the relationship between female microbiomes and competitive traits relatively understudied. Sexual experience throughout an individual’s lifetime may also be reflected in their reproductive microbiome, as age was positively correlated with cloacal microbiome richness in female tree swallows (*Tachycineta bicolor)* (Hernandez et al., 2021). Though directionality can be challenging to disentangle from observational studies, these results nevertheless suggest an important relationship between competitive phenotypes and microbial diversity. Because there are inter- and intraspecific differences in the degree of sexual selection, it is important to study microbiome-fitness dynamics among multiple species and between both sexes.

Here, we conduct an exploratory study on the reproductive microbiomes of two socially polyandrous shorebirds, *Jacana spinosa* (Northern Jacana) and *J. jacana* (Wattled Jacana). Both *J. spinosa* and *J. jacana* are socially polyandrous, in which females mate with multiple males and are typically larger in body size and weaponry, and males mate monogamously and conduct the majority of parental care (Emlen & Wrege, 2004a; Jenni & Collier, 1972). However, *J. spinosa* has a greater degree of sexual dimorphism, higher average number of male mates, and longer sperm morphology, suggesting that sexual selection is stronger in this species (Lipshutz, 2017; Lipshutz et al., 2023). Therefore, we predicted that female *J. spinosa* would have higher microbial diversity than female *J. jacana.* Given sex differences in patterns of sexual transmission, in which females mate with multiple males, but males only mate with one female, we expected higher microbial diversity among females than males. Finally, we evaluated the relationship between microbial diversity and sexually selected traits, and predicted that females with larger competitive traits would have more diverse reproductive microbiomes.

## METHODS

### Study sites and species

*Jacana spinosa* and *J. jacana* are tropical, freshwater shorebirds, and can be found in marshes and agricultural landscapes from Mexico to Central and South America. We studied wild, free-living populations of *J. spinosa* in La Barqueta, Chiriqui, Panama (8.207N, 82.579W) and *J. jacana* between Las Guabas (8.384N, 80.447) and Chepo, Panama (9.166N, 79.122W). Our fieldwork took place during the rainy season from May 16 to July 1, 2018, when breeding is most prevalent in jacanas, though jacanas may breed all year when freshwater is available (Emlen & Wrege, 2004b). We focused on reproductively active adults, including territorial females and their male mates. Male jacanas breed asynchronously and cycle between two breeding stages: courtship, when males actively copulate with their female mate, or parenting, when male are incubating eggs, brooding, or foraging with chicks (Lipshutz & Rosvall, 2020). We confirmed these breeding stages with behavioral observations of copulation, and the presence of nests and/or brood patches.

### Sample collection

In total, we sampled 39 individuals: 11 *J. spinosa* females, 12 *J. spinosa* males (five courting, seven incubating), 9 *J. jacana* females, and 7 *J. jacana* males (four courting, three incubating). These individuals were collected for a study on the neurogenomic mechanisms of social polyandry, which involved terminal collection of brain and peripheral tissues. Birds were euthanized with an air rifle, followed by an overdose of isoflurane anesthetic. These sample sizes are limited, stemming in part from the logistical challenges of tropical fieldwork in wetland habitats. Nonetheless, these samples are a valuable contribution, as the microbial communities of this avian family, Jacanidae, have not yet been described.

In birds, most research focuses on the gut microbiome (Sun et al., 2022), which is typically isolated non-lethally from fecal or cloacal samples (Berlow et al., 2020). However, the cloacal cavity is an endpoint for intestinal, urinary, and reproductive functions. Thus, cloacal samples may also reflect the reproductive microbiome, and we use the term cloacal microbiome to represent the reproductive microbiome. To collect the cloacal sample, we cleaned the outside of the cloaca with an alcohol wipe and eased a sterile swab (Puritan 25–3316-U Ultra Flocked Swab, USA) into the cloaca. Once fully inserted, we gently turned the swab for 3-5 s. Swabs were stored in 500uL of RNAlater (Invitrogen; Carlsbad, CA USA), flash frozen on dry ice, and later stored at -80°C until DNA extractions.

We measured competitive phenotypes that have previously been associated with territoriality and reproductive success in *J. jacana* (Emlen & Wrege, 2004b). We measured body mass to the nearest gram using a Pesola scale, and wing spurs to the nearest millimeter using calipers. An analysis of testosterone levels in circulation has been published for these same *J. spinosa* individuals from both sexes (Lipshutz & Rosvall, 2020) and of sperm morphology for these same males from both species (Lipshutz et al., 2023).

### DNA extraction, PCR, and sequencing

We extracted DNA from whole swabs using the DNeasy PowerSoil DNA isolation kit (Qiagen, Inc.) with minor modifications to the manufacturer’s recommended protocol (Vo & Jedlicka, 2014). We pipetted off the RNAlater supernatant from the swabs prior to DNA extraction. In addition to Solution C1, we added 10μl of Proteinase K for the tissue lysing step. The two solutions (Solutions C2 and C3) which precipitate non-DNA substances were combined (Berlow et al., 2022). Final DNA was eluted from each spin column twice with the same 100 μl of Solution C6. DNA extracts were frozen at -30°C until PCR.

PCR and sequencing follow the methods described in (Houtz et al., 2023). We amplified the V4 region of the 16S rRNA gene using the primers 515F and 806R with Illumina adapters added. We ran 10 μl PCR reactions in triplicate for each sample and included 5 μl of 2× Platinum Hot Start Master Mix (Invitrogen, Waltham, MA, USA), 0.5 μl of 10 μM primers, 3 μl of nuclease free water, and 1 μl of template DNA. Cycling conditions were 3 min at 94°C followed by 35 cycles of 94°C for 45 s, 50°C for 60 s, and 72°C for 90 s before a final extension at 72°C for 10 min. We pooled the three replicate reactions for each sample and ran a 1% agarose gel to confirm that amplification was successful for the V4 region of the 16S rRNA gene (∼ 350 bp). Each PCR run included negative controls (nuclease-free water in place of template DNA but were not sequenced). We also extracted, amplified, and sequenced 4 negative kit reagent controls. We submitted our final pooled PCR products to the Cornell Biotechnology Resource Center for quantification, normalization, library preparation, and sequencing on a single Illumina MiSeq run (2 x 250 bp).

### Cloacal microbiome bioinformatics

We used QIIME2 (Quantitative Insights Into Microbial Ecology; v 2024.5) to process demultiplexed paired-end forward and reverse sequences as FASTQ files (Bolyen et al., 2019). We used the DADA2 plugin to remove primers, truncate to 180 bp, and create amplicon sequence variants (ASVs). We assigned taxonomy to the ASVs by fitting a naive-Bayes classifier trained on the Silva 138 database using the sk-learn classifier (Yilmaz et al., 2014). The phylogeny plugin was applied to construct a rooted phylogenetic tree by employing FastTree and MAFFT. We removed singletons, chloroplasts, mitochondria, eukaryotes, archaea, and any ASVs unassigned to a bacterial phylum.

We exported the filtered ASV table, taxonomy, and phylogenetic tree artifacts from QIIME2 into R v 4.3.1 for all subsequent analyses. Using the ‘phyloseq’ package v 1.46.0, we combined the ASV table, sample metadata, taxonomy table, and phylogenetic tree into a phyloseq object (McMurdie & Holmes, 2013). We used the ‘decontam’ package v 1.22.0 to remove 6 contaminant ASVs and were left with 2,323 ASVs across 39 total samples after removing negative controls (n = 4) (Davis et al., 2018).

To limit bias due to differing sequencing depths, we rarefied samples to 14,000 reads, removing 2 samples (WAF3 and WAM6) with less than 14,000 reads, leaving 2,080 ASVs across 37 samples. With the ‘phyloseq’ package, we calculated alpha diversity metrics including Chao1 index (Chao, 1984) and Shannon index (Shannon & Weaver, 1949). Faith’s Phylogenetic Diversity (PD) (Faith, 1992) was calculated using the ‘picante’ package v 1.8.2 (Kembel et al., 2010). Chao1 index measures the richness of the samples, Shannon’s index considers the richness and evenness of the samples, and Faith’s PD assigns the diversity score based on how related the species are to each other from our phylogenetic tree.

For bacterial beta diversity, we calculated Bray-Curtis dissimilarity (richness and abundance of the ASVs in the communities; (Sørensen, 1948) and Jaccard distance (ASV richness only; (Jaccard, 1908). All samples were processed to remove exceptionally rare taxa that could disproportionately influence beta diversity metrics. ASVs were excluded if they had fewer than 50 reads in total across all samples. Bray-Curtis dissimilarity and Jaccard matrices were calculated using reads rarefied to 14,000 as described for alpha diversity metrics. After prevalence filtering, 353 ASVs were retained from rarefied data.

### Statistical analysis

All statistical analyses were run in R v 4.3.1. We used the package ‘lme4’ v 1.1-35.3 (Bates et al., 2015) to run linear mixed effect models with each microbial alpha diversity. The package ‘DHARMA’ v 0.4.6 was used to carry out model diagnostics (Hartig, 2018). To test for changes in microbial beta diversity, we conducted a separate permutational multivariate analysis of variation (PERMANOVA) with each distance matrix (Bray-Curtis or Jaccard) as the response variable to statistically partition the sources of variation in microbial community structure with the *adonis2* function in the ‘vegan’ package v v 2.6-6.1 (Oksanen et al., 2022), with the ‘by’ parameter set for ‘margin’ to account for marginal effects of the tested variables. An assumption of the adonis test is that groups have homogeneity of variance. The dispersion of the groups outlined in each section below was checked for homogeneity of variance using the *Betadisper* and *permutest* functions from the ‘vegan’ package. All figures were created using the ‘ggplot2’ package v 3.5.1 (Wickham, 2016) and cartoons were added in PowerPoint.

We used a Pearson correlation to compare competitive traits, log testosterone levels, and alpha diversity metrics using the cor.test function in R. We analyzed correlation matrices using the cor.mtest function and visualized them with the package corrplot (Wei & Simko, 2024). We conducted a similarity percentage (SIMPER) analysis to identify the average contribution of each bacterial genus to the Bray–Curtis dissimilarity (beta diversity) of the significant species comparison in PAST (v 4.15), with a 70% cutoff for low contributions to dissimilarity. Beta diversity ordinations for Bray-Curtis and Jaccard were visualized with principal component analyses (PCoA) calculated with the ‘phyloseq’ package. We only tested for relationships between bacterial taxa in female *J. spinosa* because we found relationships between testosterone and alpha diversity in female *J. spinosa*. We used MaAsLin2 (Microbiome Multivariable Associations with Linear Models; (Mallick et al., 2021) to identify differentially-abundant taxa with testosterone in female *J. spinosa* as the fixed effect with the following functions: analysis_method = “CPLM” (Compound Poisson Linear Model), transform = “AST” (Arcsine Transformation), and correction = “BH” (Benjamini-Hochberg or False Discovery Rate for multiple comparisons correction). We used the rarefied ASV table as the input for the MaAsLin2 analysis, as MaAsLin2 based on rarefied data produces the most consistent results across datasets (Nearing et al., 2022).

## RESULTS

### Sequencing results

Before filtering, there were 1,254,953 total reads and 2,445 unique ASVs across 43 samples (n = 39 cloacal samples, n = 4 negative controls). Mean reads per sample were 29,185 with samples ranging from 257 to 53,578 reads. After filtering (i.e., removal of removed singletons, chloroplasts, mitochondria, eukaryotes, archaea, and any ASVs unassigned to a bacterial phylum), we retained 1,222,235 reads and 2,329 unique ASVs that were used to calculate gut microbial alpha and beta diversity metrics. Mean reads per sample were 28,424.1 with samples ranging from 221 to 53,022 reads.

### Species and sex differences

#### Microbial alpha diversity

We did not find a significant relationship with alpha diversity and species, sex, nor their interaction for all three diversity metrics including Chao1 index (sex*species: β = 27.63, CI = -73.04 – 128.29, P = 0.58, Supplementary Material Table S1, Supplementary Material Figure S1A), Shannon index (sex*species: β = -0.34, CI = -1.30 – 0.62, P = 0.48, Supplementary Material Table S2, Supplementary Material Figure S1B), and Faith’s PD (sex*species: β = 0.83, CI = -7.06-8.73, P = 0.83, Supplementary Material Table S3, Supplementary Material Figure S1C).

#### Microbial beta diversity

For Bray-Curtis dissimilarity, we found a significant relationship with beta diversity and species (F = 1.73, P = 0.04), but not for sex (F = 1.30, P = 0.15) nor their interaction (sex*species: F = 1.15, P = 0.25) (Supplementary Material Figure S2). There was no significant effect of species (F = 0.37, P = 0.55) or sex (F = 3.08, P = 0.09) on Bray-Curtis beta dispersion. For Jaccard distance, we also found a significant relationship with beta diversity and species (F = 1.29, P = 0.03), but not for sex (F = 1.07, P = 0.24) nor their interaction (sex*species: F = 1.08, P = 0.21) (**Figure 1**). There was no significant effect of species (F = 0.03, P = 0.84) or sex (F = 2.07, P = 0.16) on Jaccard beta dispersion.

**Figure 1.**
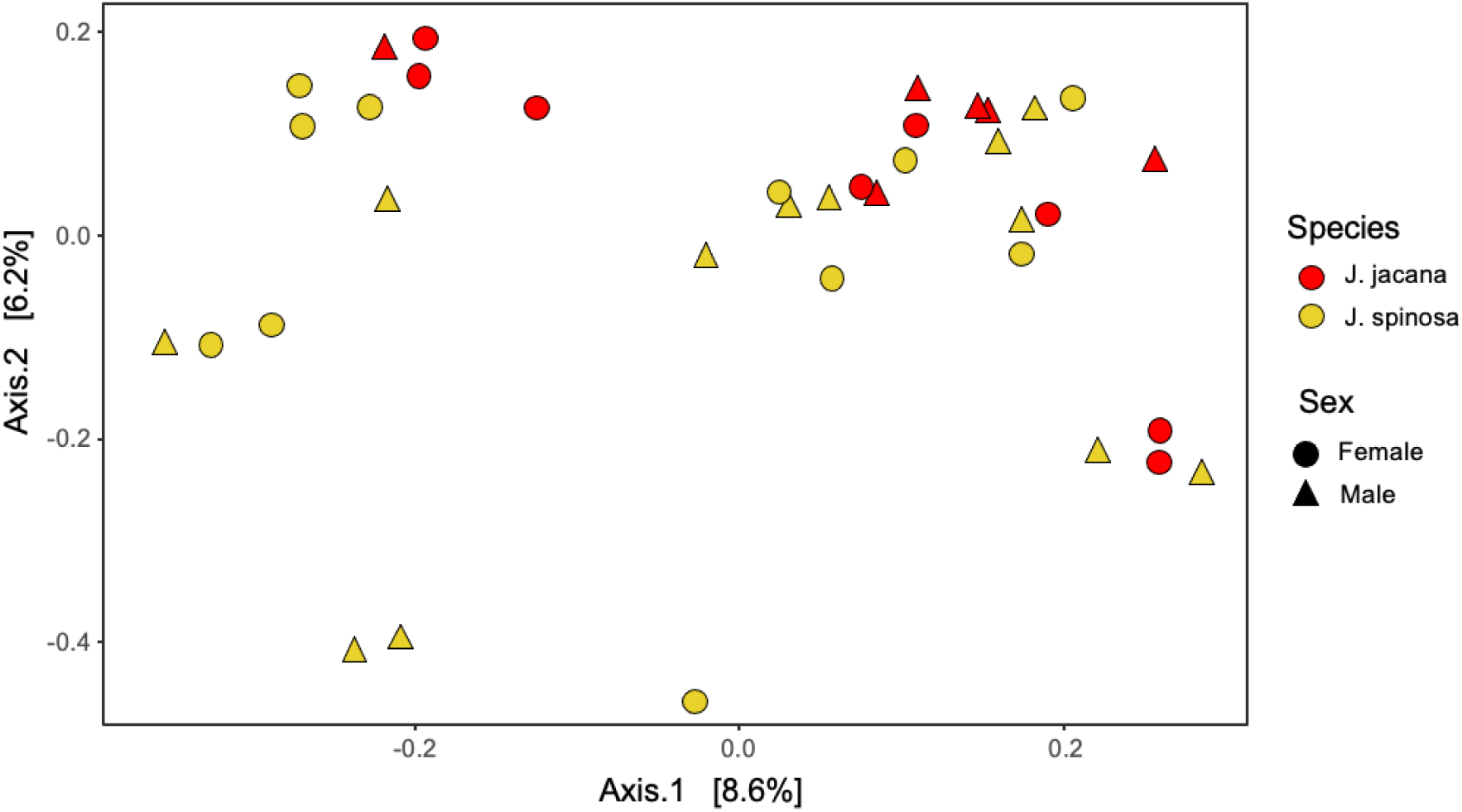
Principal component analysis (PCoA) of Jaccard distance (beta diversity) shows species differences between *Jacana spinosa* (Northern Jacana) in yellow and *J. jacana* (Wattled Jacana) in red, but not sex differences between females as circles and males as triangles.

The SIMPER analysis revealed that over 80% of the dissimilarity between *J. jacana* and *J. spinosa* can be attributed to changes in the relative abundances of 3 bacterial genera. A higher relative abundance of an unassigned genus from class Bacilli in *J. spinosa* explained 35.2% of the average dissimilarity between the two species. A higher relative abundance of *Enterococcus* and *Exiguobacterium* in *J. jacana* explained 24.23% and 22.75% of the dissimilarity between species, respectively (Supplementary Material Table S4). Other notable species of lower relative abundance across both species and sexes include *Carnobacterium maltaromaticum* and *Variovorax paradoxus* (see Supplementary Material Figure S3 for relative abundance taxa barplots).

### Alpha diversity and bacterial taxa correlate with competitive phenotypes

In female *J. spinosa*, testosterone levels positively correlated with chao1 (r = 0.74, t = 3.13, df = 8, p = 0.014) and Faith’s PD alpha diversity (r = 0.66, t = 2.51, df = 8, p = 0.036) (Figure 2A). Testosterone in female *J. spinosa* also correlated with wing spur length, a weapon (r = 0.76, t = 3.36, df = 8, p = 0.01) (Figure 2A).

**Figure 2.**
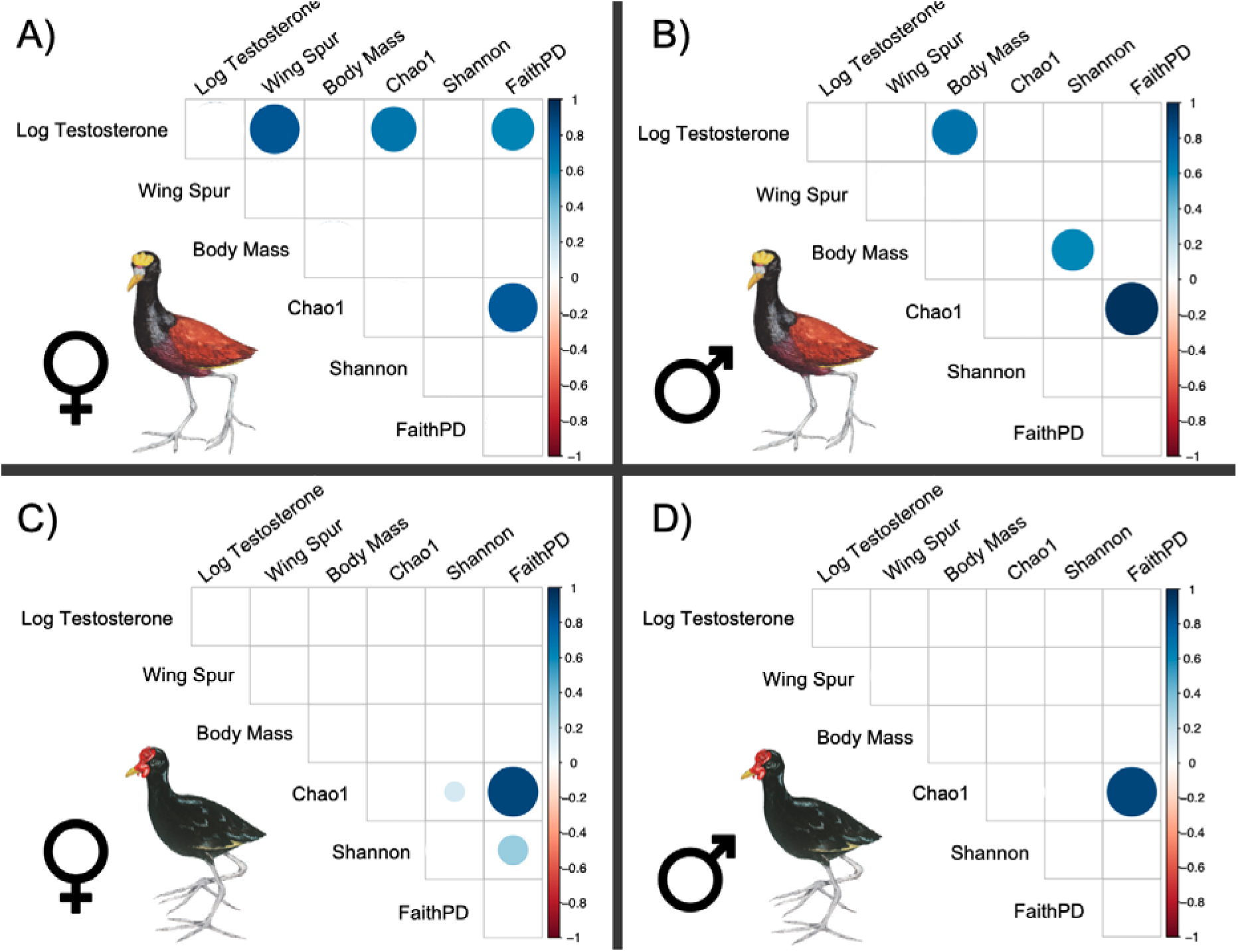
Correlation matrices for competitive traits and alpha diversity metrics for A) female *Jacana spinosa*, B) male *J. spinosa*, C) female *J. jacana*, and D) male *J. jacana*. The size and color intensity of the blue circle indicates a more strongly positive correlation.

In male *J. spinosa*, body mass correlated with Shannon diversity (r = 0.61, t = 2.42, df = 10, p = 0.036) and with testosterone levels (r = 0.69, t = 3.02, df = 10, p = 0.013) (Figure 2B). A previous study of these same individuals found that courting males had both higher testosterone and larger body mass than parenting males, so this pattern is likely driven by the combination of males in both breeding stages (Lipshutz & Rosvall, 2020). We were unable to study male breeding stages separately due to small sample sizes. In *J. jacana*, competitive traits did not correlate with alpha diversity metrics, for either sex, though several diversity metrics did correlate with one another (Figure 2C, 2D).

The MaAsLin2 analysis found significant correlations between testosterone and 40 differentially-abundant ASVs in female *J. spinosa* (Supplementary Material Table S5). Testosterone levels were positively correlated with 28 ASVs and negatively correlated with 12 ASVs. Of note, testosterone levels were negatively correlated with two ASVs assigned as *Exiguobacterium* (ASV1: coef = -477.48, p < 0.001; ASV2: coef = -476.46, p < 0.001) and 1 ASV assigned to *Enterococcus gallinarum* (coef = -439.01, p < 0.001). Testosterone levels in female *J. spinosa* were positively correlated with an ASV assigned to *Variovorax paradoxus* (coef = 36.12, p < 0.001), among others (Supplementary Material Table S5).

## DISCUSSION

We hypothesized that social polyandry would predict cloacal microbial diversity, such that it would be higher in *J. spinosa* and in females. We found no relationship between alpha diversity and species or sex, but we did find that beta diversity significantly differed between species. We also hypothesized that microbial diversity would be associated with competitive traits. We found a positive relationship between microbial alpha diversity and testosterone and weaponry for female *J. spinosa*. Based on our alpha diversity results, we also tested for relationships between testosterone and relative abundance of bacterial genera in female *J. spinosa*. We found significant correlations between testosterone and 40 ASVs in female *J. spinosa*.

### Microbial alpha diversity is associated with competitive traits

We show that female *J. spinosa* with higher microbial alpha diversity also had higher testosterone levels and longer wing spurs. A similar pattern was found in male rufous collared sparrows (*Zonotrichia capensis*), in which higher testosterone concentrations were associated with greater diversity of cloacal bacteria (Escallón et al., 2016). This increase in diversity may be due to testosterone’s role in promoting sexual behaviors, as male testosterone levels are higher in polygynous mating systems (Garamszegi et al., 2005) and associated with more frequent extra-pair fertilizations (Raouf et al., 1997). Jacanas defend territories containing multiple male mates, and females with the largest competitive phenotypes have higher reproductive success (Emlen & Wrege, 2004b). However, we do not know whether females with higher alpha diversity had more sexual partners, as we did not measure this in the field.

An alternative, but not mutually exclusive explanation, is that higher testosterone suppresses immune function, facilitating the colonization of additional microbes in the reproductive tract. This potential trade-off between competitive phenotypes, sexual transmission, and immune function may explain the relationship between testosterone concentrations and the relative abundance of *Chlamydia*, a potentially pathogenic bacteria, in rufous collared sparrows (Escallón et al., 2016). In male *J. spinosa*, testosterone levels correlated with body mass, a trait likely confounded by male breeding stage, as parenting males have smaller body mass and lower testosterone than courting males (Lipshutz & Rosvall, 2020). Although we are unsure why the relationship between alpha diversity and competitive traits was only significant in *J. spinosa*, one explanation is that sexual selection is strongest in this species. *J. spinosa* females have a greater degree of polyandry, and more extreme female-biased sexual dimorphism than *J. ajcana* females (Lipshutz, 2017). Future work should test whether female jacanas with more mating partners receive more diverse microbes via sexual transmission, to directly test this relationship.

### Cloacal beta diversity differs by species but not sex

We predicted that *J. spinosa* would have higher microbial alpha diversity than *J. jacana*, due to species differences in the strength of sexual selection (Lipshutz, 2017; Lipshutz et al., 2023). We also expected higher microbial diversity among females than males, because of their socially polyandrous mating systems. However, we did not find significant differences in microbial alpha diversity between species nor between sexes. We found a significant effect of species on cloacal microbial beta diversity, but no effect of sex. Although gut microbial alpha diversity did not differ between sexes or species, beta diversity differences suggest that *J. spinosa* and *J. jacana* may differ in their gut microbial community composition while maintaining similar alpha diversity levels. Though other studies have found no sex differences in microbial diversity e.g. (Gongora et al., 2021), future work should re-examine these patterns with larger sample sizes.

*J. spinosa* and *J. jacana* may differ in their microbial compositions due to geographic isolation or dietary strategy. Previous ecological niche modeling found significant differences in habitat suitability between the species, with suitable habitats for *J. spinosa* being wetter and warmer (Miller et al., 2014). The two species do hybridize where they overlap (Lipshutz et al., 2019), but our samples are from individuals outside the hybrid zone. Observations from the field suggest that both species eat seeds and aquatic vegetation, as well as insects, aquatic invertebrates, and small fish and amphibians (Jenni & Kirwan, 2020). We found the presence of *Camobacterium maltaromaticum*, a bacterial species found in freshwater habitats (Tang et al., 2023) and animal prey of jacanas including fish (Iskandar et al., 2017), and insects (Leisner et al., 2007). Likewise, another bacterial genus found in both species, *Variovorax paradoxus*, is also found in freshwater ecosystems (Dul’tseva et al., 2012). Future research should characterize the diets of both species via diet metabarcoding of fecal samples and investigate potential relationships with the microbiome.

Sexual transmission of bacteria in birds may be asymmetrical, as males copulate into the cloaca of the female (Kulkarni & Heeb, 2007); however, how these patterns change in polyandrous vs. polygynous systems is still an open question. Within each species, we found no difference in microbial beta diversity between males and females. At the population level, sex differences in the diversity of reproductive microbiomes may be eroded by increasing sexual transmission (Rowe et al., 2020). Males and females may homogenize their gut microbial compositions through copulations. Kittiwakes share a similar cloacal microbial composition with their mates, but after inseminations are experimentally blocked, the cloacal communities of mates became increasingly dissimilar (White et al., 2010). Transmission of cloacal bacteria between males and females is likely limited to the breeding season (Escallon et al., 2019), when there is direct cloacal contact between individuals. Thus, the year round breeding season of jacanas (Emlen & Wrege, 2004b) allows for continuous intersexual transmission of cloacal microbiota.

### Conclusions

Though this was an exploratory study, the patterns we uncovered between microbial diversity and competitive traits pose exciting new directions for understanding female-driven dynamics shaping reproductive microbiomes in wild birds. Promising research directions should include experimental manipulations of the reproductive microbiome via antibiotics or probiotic supplementation to investigate causal links between the microbiome and competitive phenotypes.

## ACKNOWLEDGEMENTS

We thank Elizabeth Derryberry for conceptual and logistical support. Evan Buck, Clara Howell, and Jose Alejandro Ramirez Silva for assistance in the field. We thank Matt Fuxjager and M. Miles for assistance importing samples. We thank Daniel Ernesto Buitrago for assistance translating the abstract into Spanish. We thank anonymous reviewers whose feedback improved the manuscript.

## Ethics statement

Scientific collection in Panama was done with permission from landowners and prior approval of MiAmbiente, Panama’s environmental authority (permit SE/A-17-18). Methods were evaluated and approved by the Institutional Animal Care and Use Committee (IACUC) of the Smithsonian Tropical Research Institute (IACUC permit 2018-0116-2021) and the University of Tennessee (IACUC permit 2573).

## Author Contributions

Study conception: S.E.L. and M.B.; data collection: S.E.L; labwork: M.B. and J.L.H; computation: K.A.A., J.L.H; data analysis: K.A.A., J.L.H, S.E.L.; resources and funding acquisition: S.E.L. and J.L.H; writing: S.E.L., K.A.A., J.L.H.

## Data Availability

All files necessary for replicating our processing and analysis, including Custom QIIME2, R scripts and data files, are available on GitHub (https://github.com/slipshut/Jacana_Microbiome). Sequences are available at the National Center for Biotechnology Information (NCBI) under BioProject PRJNA1204956.

## Supplemental Material

**Supplementary Material Figure S1.**
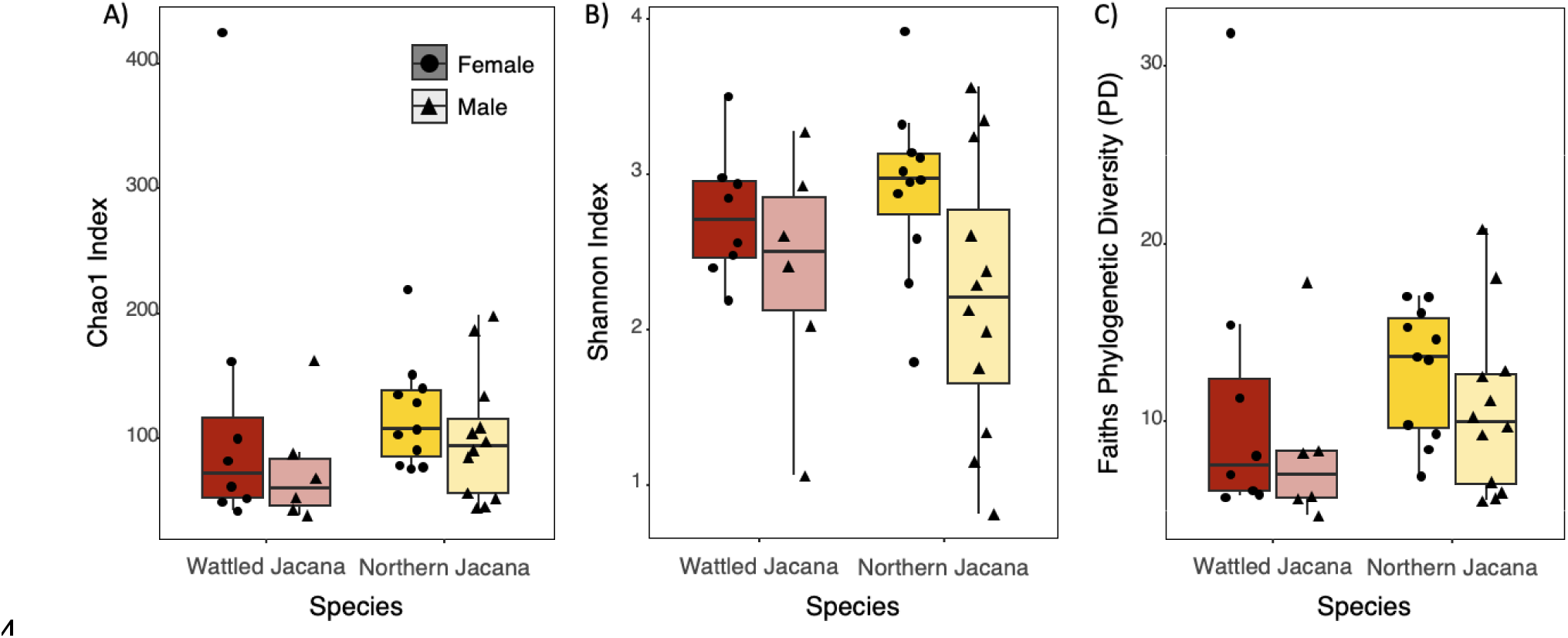
Alpha diversity between species and sexes in A) Chao1 index, B) Shannon index, and C) Faith’s Phylogenetic Diversity (PD). Red indicates *Jacana jacana* (Wattled Jacanas) and yellow indicates *J. spinosa* (Northern Jacanas). Darker colors and circles indicate females, and lighter colors and triangles indicate males. There were no significant differences between species, nor sexes.

**Supplementary Material Figure S2:**
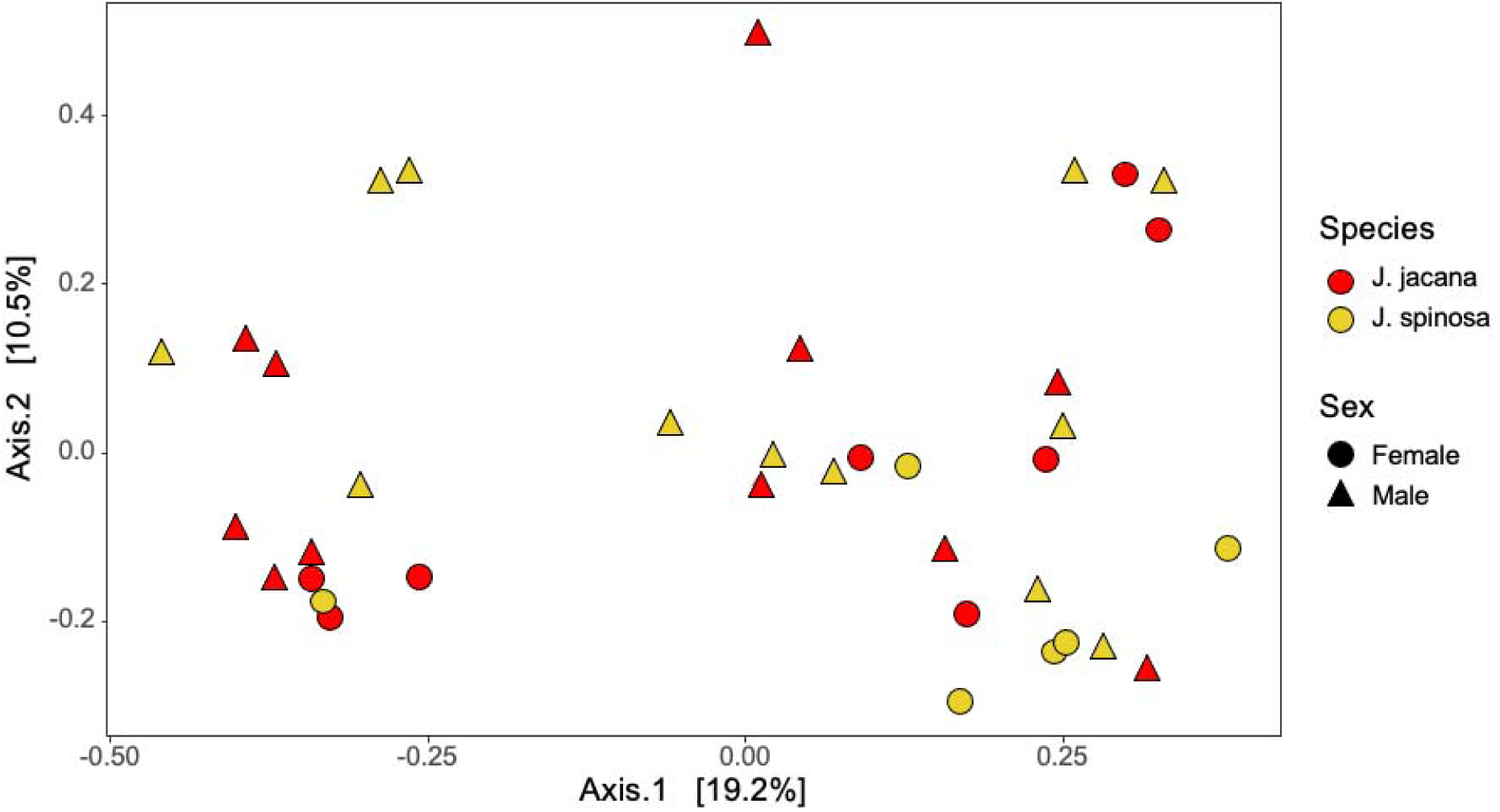
Principal coordinates analysis (PCoA) of Bray-Curtis dissimilarity of jacana gut microbiomes by species and sex. Red indicates *Jacana jacana* (Wattled Jacanas) and yellow indicates *J. spinosa* (Northern Jacanas). Circles represent females and triangles represent males.

**Supplementary Material Figure S3:**
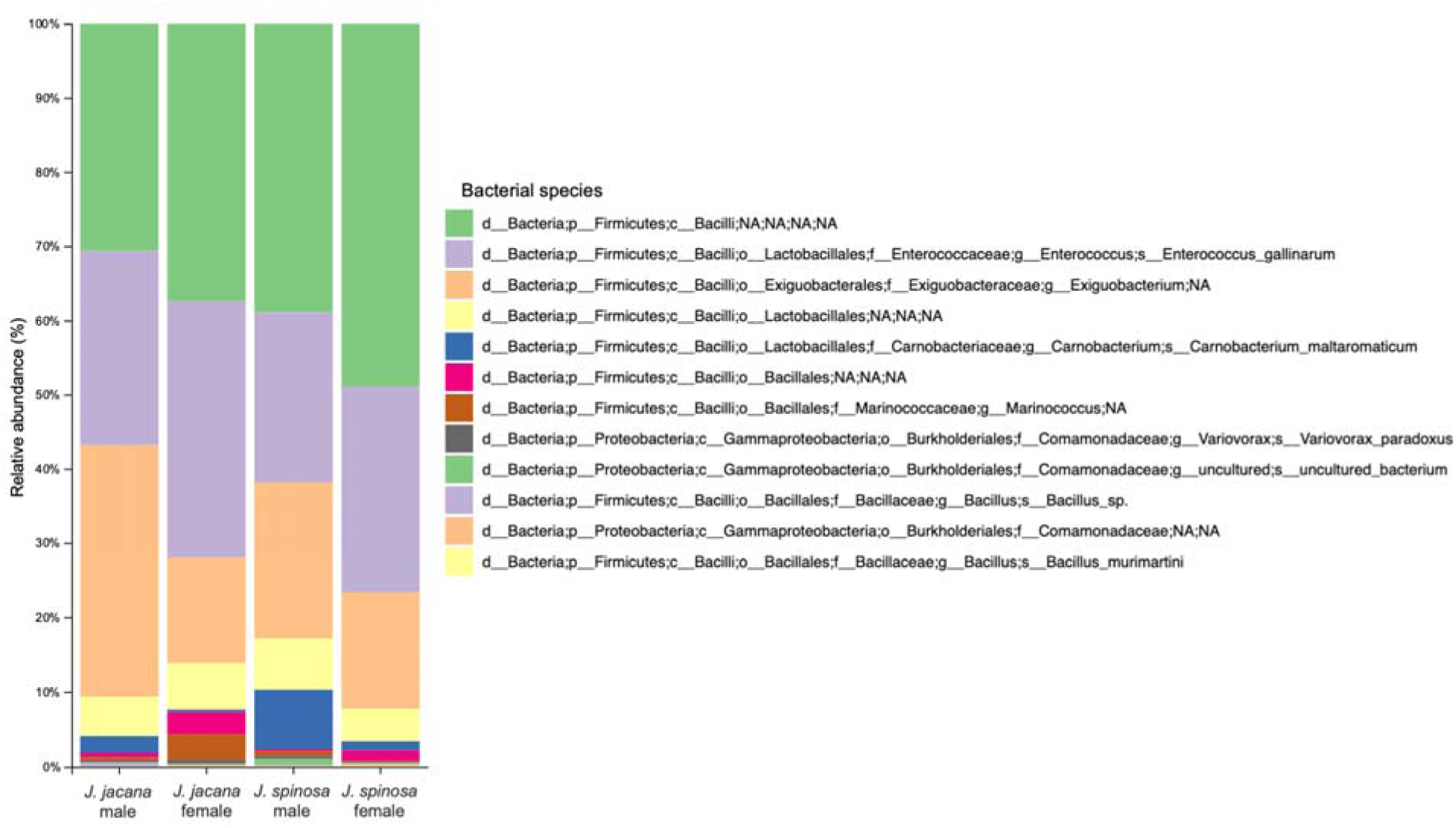
Relative abundance of bacterial species of jacana gut microbiomes by species and sex including female and male *Jacana jacana* (Wattled Jacanas) and *J. spinosa* (Northern Jacanas).

**Supplementary Material Table S1.**
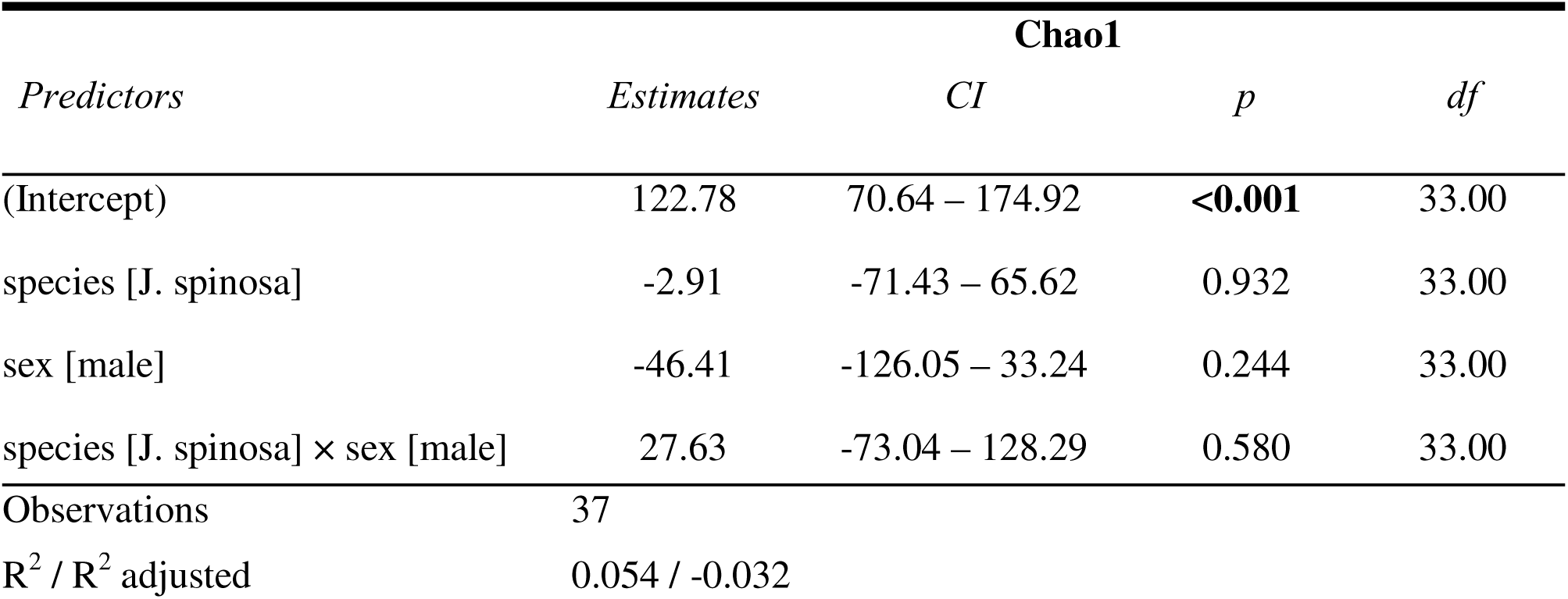
Linear model testing species x sex interaction on Chao1 index.

**Supplementary Material Table S2.**
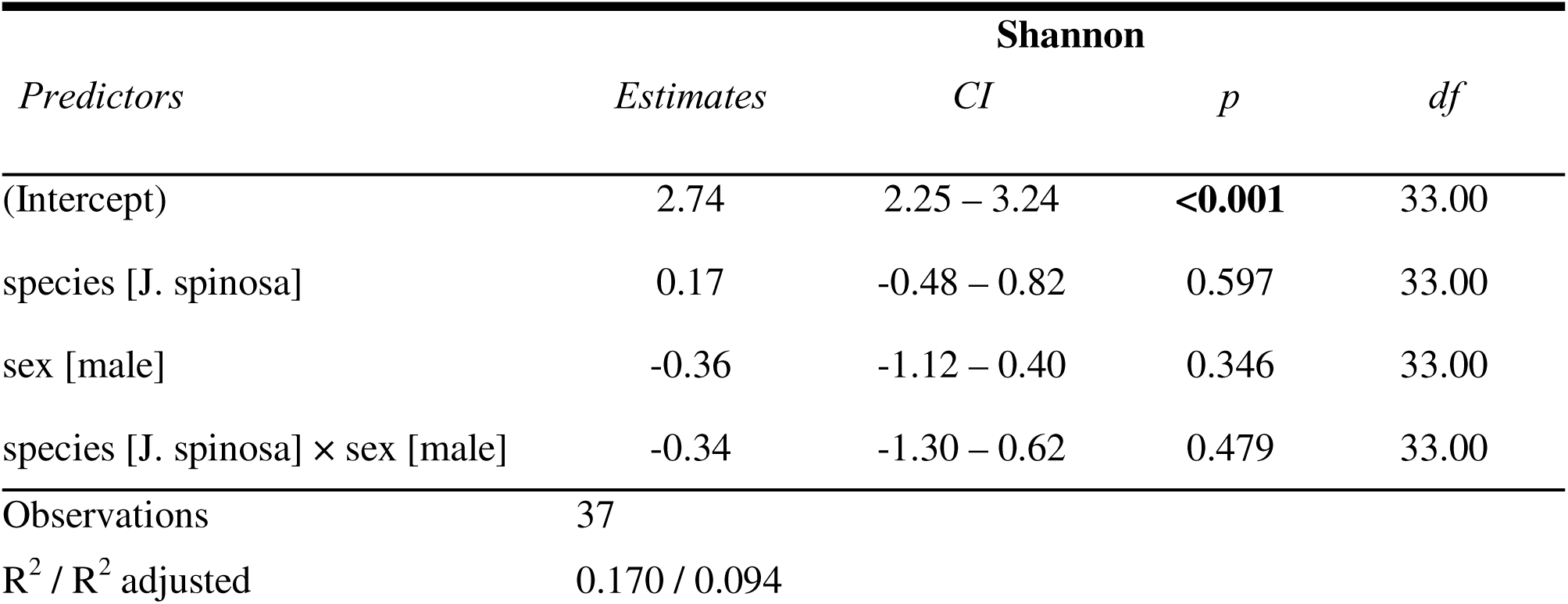
Linear model testing species x sex interaction on Shannon index.

**Supplementary Material Table S3.**
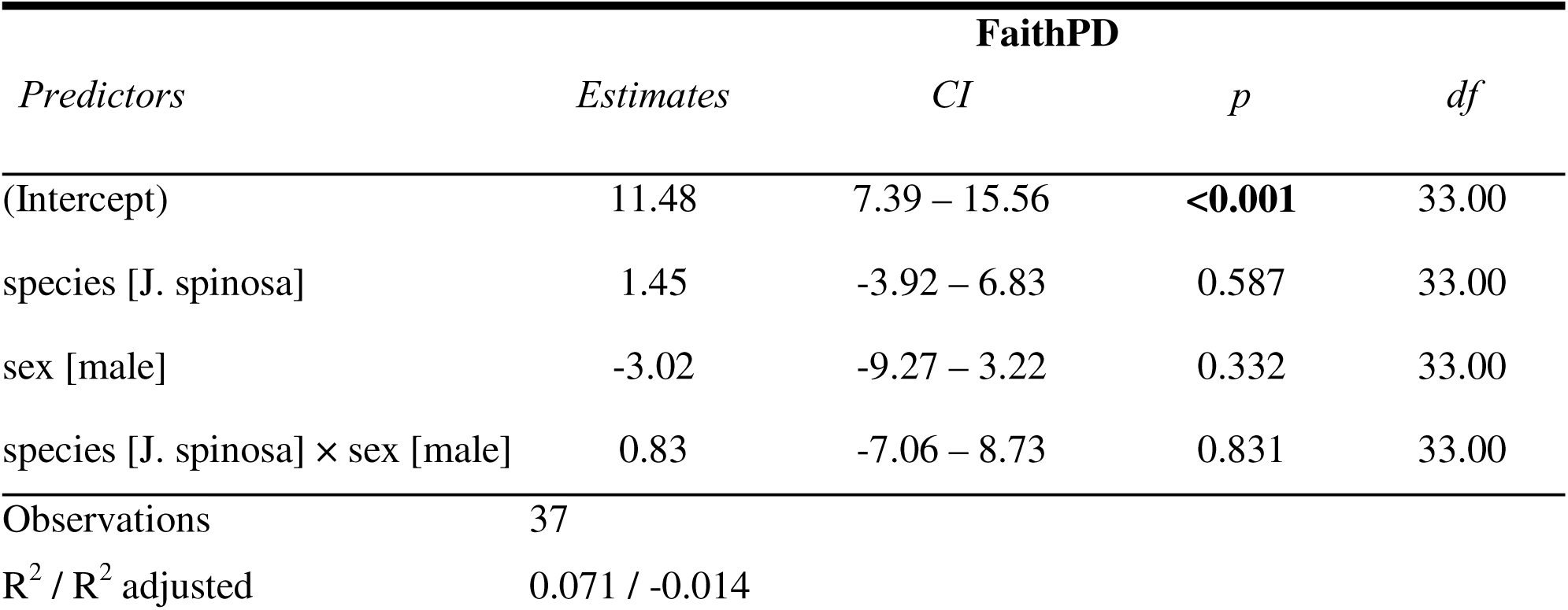
Linear model testing species x sex interaction on Faith’s Phylogenetic Diversity (PD).

**Supplementary Material Table S4.**
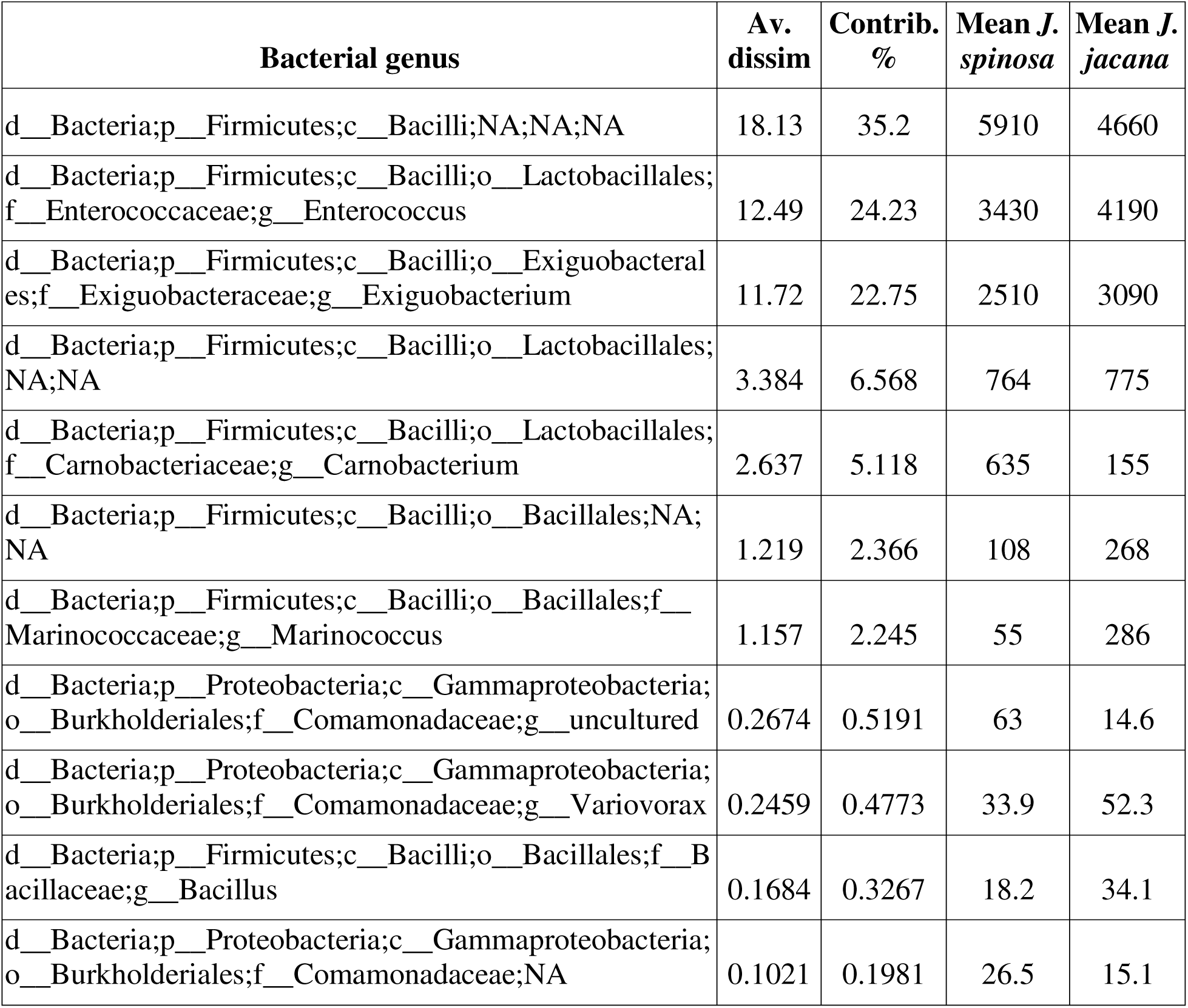
Similarity percentage (SIMPER) analysis to identify the average contribution of each bacterial genus to the Bray-Curtis dissimilarity between species.

**Supplementary Material Table S5.**
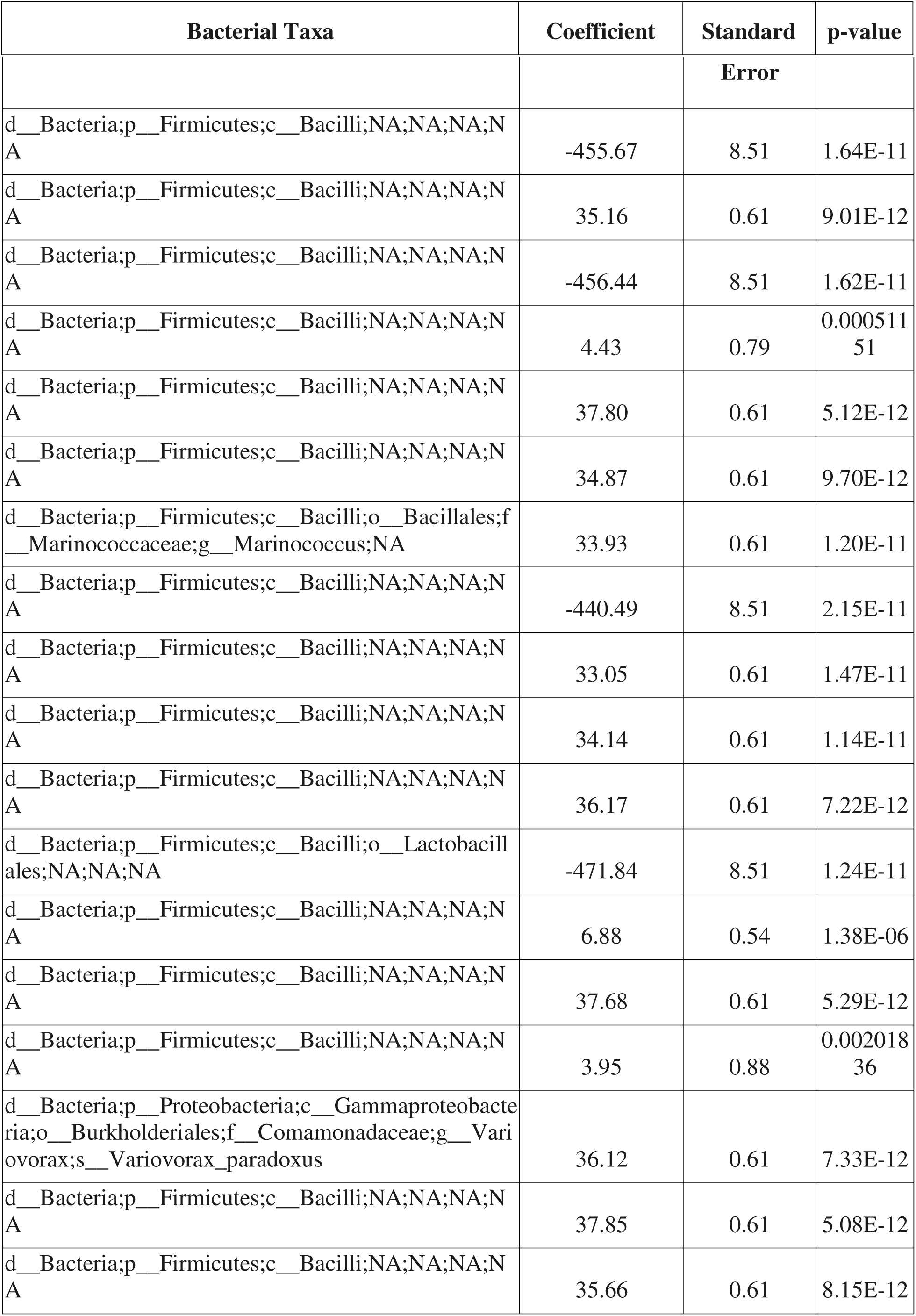
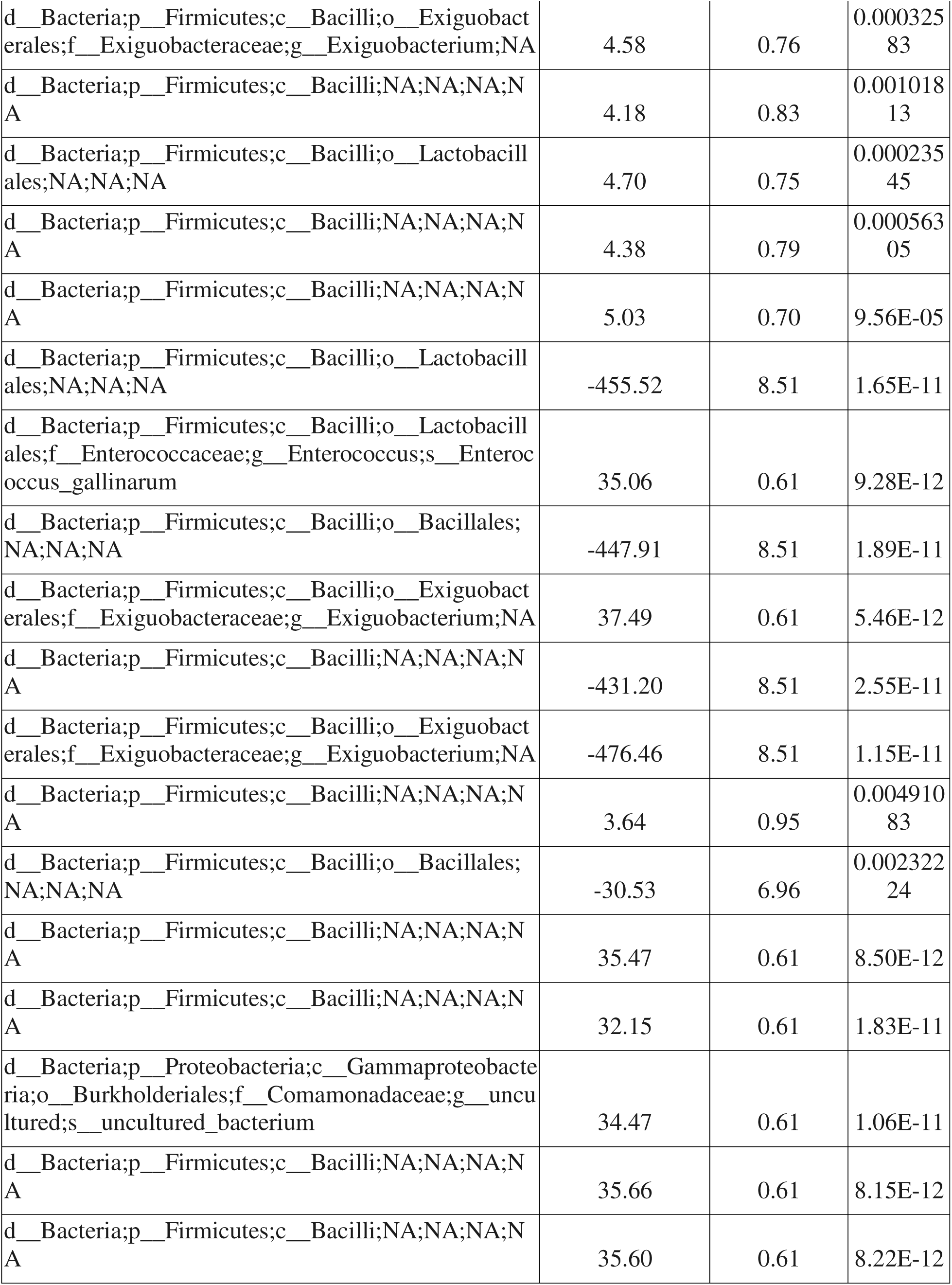
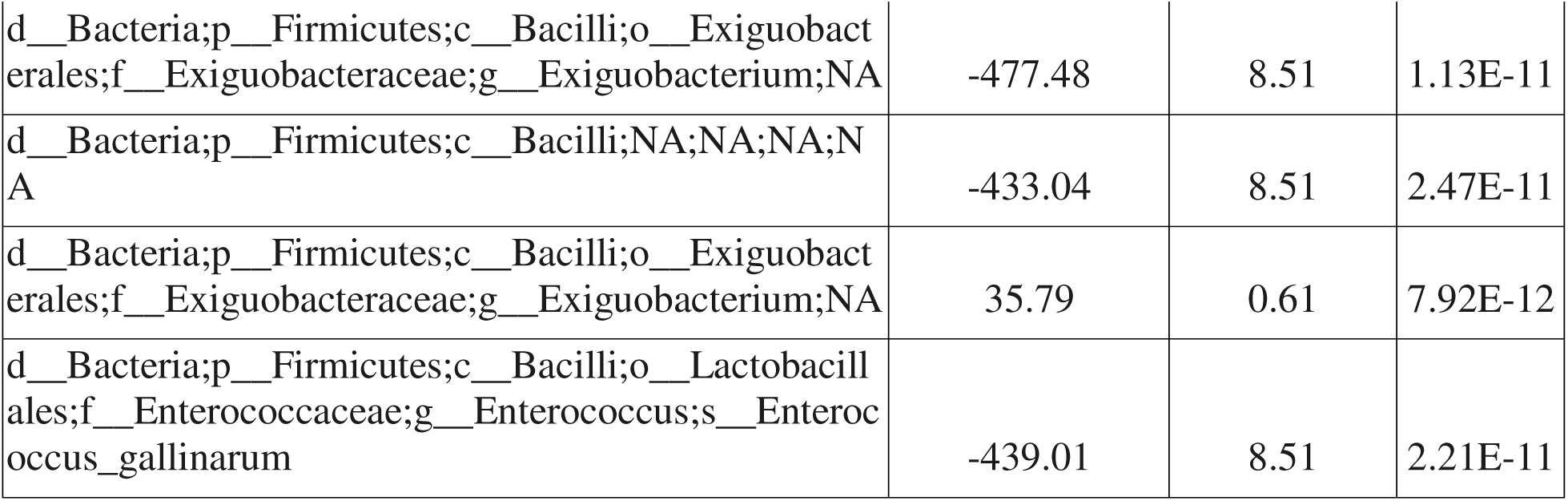
Differentially-abundant bacterial taxa correlated with testosterone in female northern jacanas from the MaAsLin2 analysis.

