## Supplementary Material for "Reproductive microbial diversity is associated with competitive phenotypes in socially polyandrous jacanas"

**Supplemental Material**

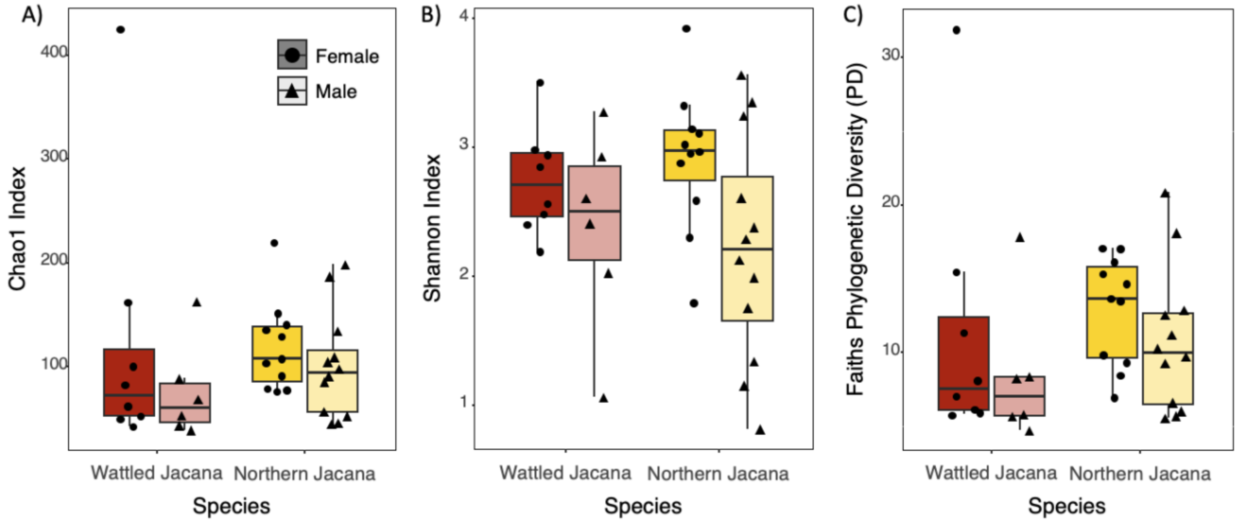


**Supplementary Material Figure S1**. Alpha diversity between species and sexes in A) Chao1 index, B) Shannon index, and C) Faith's Phylogenetic Diversity (PD). Red indicates *Jacana jacana* (Wattled Jacanas) and yellow indicates *J. spinosa* (Northern Jacanas). Darker colors and circles indicate females, and lighter colors and triangles indicate males. There were no significant differences between species, nor sexes.

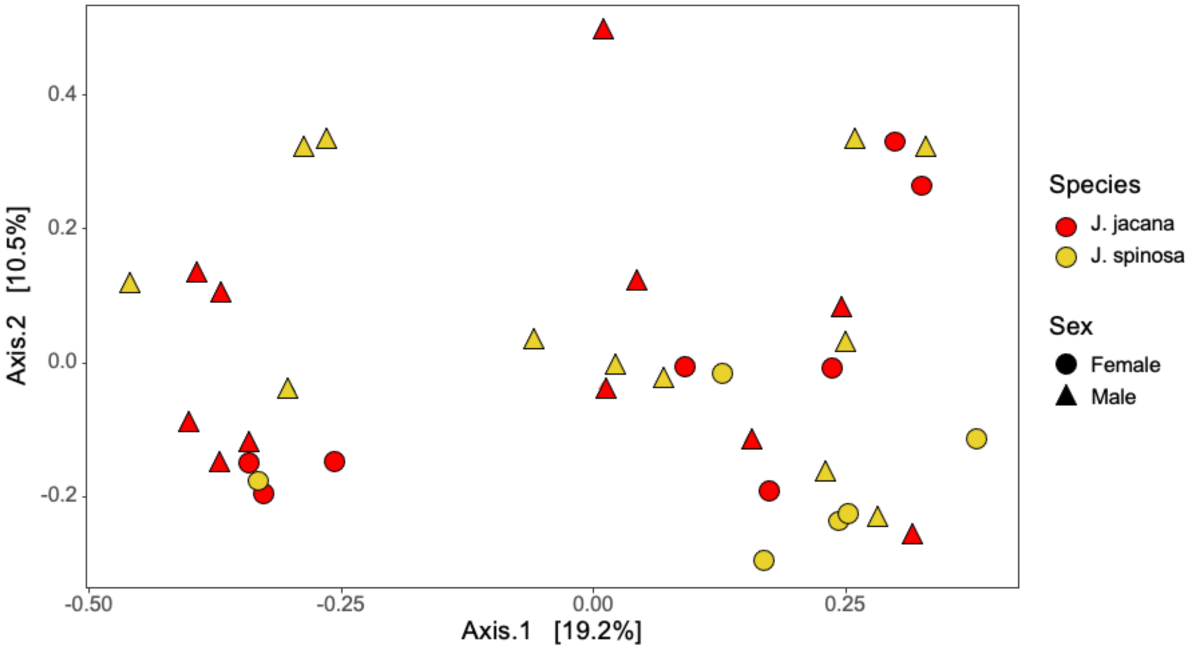


**Supplementary Material Figure S2**: Principal coordinates analysis (PCoA) of Bray-Curtis dissimilarity of jacana gut microbiomes by species and sex. Red indicates *Jacana jacana* (Wattled Jacanas) and yellow indicates *J. spinosa* (Northern Jacanas). Circles represent females and triangles represent males.

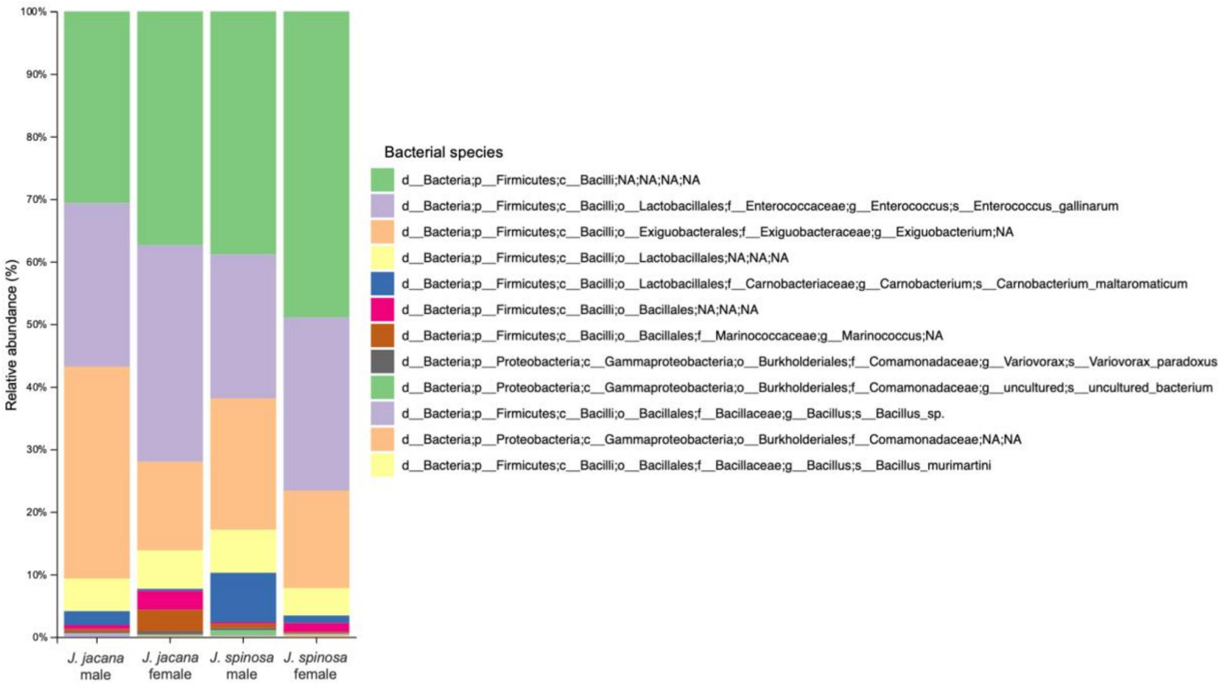


**Supplementary Material Figure S3**: Relative abundance of bacterial species of jacana gut microbiomes by species and sex including female and male *Jacana jacana* (Wattled Jacanas) and *J. spinosa* (Northern Jacanas).

**Supplementary Material Table S1.** Linear model testing species x sex interaction on Chao1 index.

| | Chao1 | | | |
|---|---|---|---|---|
| *Predictors* | *Estimates* | *CI* | *p* | *df* |
| (Intercept) | 122.78 | 70.64 – 174.92 | **<0.001** | 33.00 |
| species [J. spinosa] | -2.91 | -71.43 – 65.62 | 0.932 | 33.00 |
| sex [male] | -46.41 | -126.05 – 33.24 | 0.244 | 33.00 |
| species [J. spinosa] × sex [male] | 27.63 | -73.04 – 128.29 | 0.580 | 33.00 |
| Observations | 37 | | | |
| $R^2$ / $R^2$ adjusted | 0.054 / -0.032 | | | |

**Supplementary Material Table S2.** Linear model testing species x sex interaction on Shannon index.

| | Shannon | | | |
|---|---|---|---|---|
| *Predictors* | *Estimates* | *CI* | *p* | *df* |
| (Intercept) | 2.74 | 2.25 – 3.24 | **<0.001** | 33.00 |
| species [J. spinosa] | 0.17 | -0.48 – 0.82 | 0.597 | 33.00 |
| sex [male] | -0.36 | -1.12 – 0.40 | 0.346 | 33.00 |
| species [J. spinosa] × sex [male] | -0.34 | -1.30 – 0.62 | 0.479 | 33.00 |
| Observations | 37 | | | |
| $R^2$ / $R^2$ adjusted | 0.170 / 0.094 | | | |

**Supplementary Material Table S3.** Linear model testing species x sex interaction on Faith's Phylogenetic Diversity (PD).

| | FaithPD | | | |
|---|---|---|---|---|
| *Predictors* | *Estimates* | *CI* | *p* | *df* |
| (Intercept) | 11.48 | 7.39 – 15.56 | **<0.001** | 33.00 |
| species [J. spinosa] | 1.45 | -3.92 – 6.83 | 0.587 | 33.00 |
| sex [male] | -3.02 | -9.27 – 3.22 | 0.332 | 33.00 |
| species [J. spinosa] × sex [male] | 0.83 | -7.06 – 8.73 | 0.831 | 33.00 |
| Observations | 37 | | | |
| $R^2$ / $R^2$ adjusted | 0.071 / -0.014 | | | |

**Supplementary Material Table S4.** Similarity percentage (SIMPER) analysis to identify the average contribution of each bacterial genus to the Bray-Curtis dissimilarity between species.

| Bacterial genus | Av. dissim | Contrib. % | Mean *J. spinosa* | Mean *J. jacana* |
|---|---|---|---|---|
| d__Bacteria;p__Firmicutes;c__Bacilli;NA;NA;NA | 18.13 | 35.2 | 5910 | 4660 |
| d__Bacteria;p__Firmicutes;c__Bacilli;o__Lactobacillales;f__Enterococcaceae;g__Enterococcus | 12.49 | 24.23 | 3430 | 4190 |
| d__Bacteria;p__Firmicutes;c__Bacilli;o__Exiguobacterales;f__Exiguobacteraceae;g__Exiguobacterium | 11.72 | 22.75 | 2510 | 3090 |
| d__Bacteria;p__Firmicutes;c__Bacilli;o__Lactobacillales;NA;NA | 3.384 | 6.568 | 764 | 775 |
| d__Bacteria;p__Firmicutes;c__Bacilli;o__Lactobacillales;f__Carnobacteriaceae;g__Carnobacterium | 2.637 | 5.118 | 635 | 155 |
| d__Bacteria;p__Firmicutes;c__Bacilli;o__Bacillales;NA;NA | 1.219 | 2.366 | 108 | 268 |
| d__Bacteria;p__Firmicutes;c__Bacilli;o__Bacillales;f__Marinococcaceae;g__Marinococcus | 1.157 | 2.245 | 55 | 286 |
| d__Bacteria;p__Proteobacteria;c__Gammaproteobacteria;o__Burkholderiales;f__Comamonadaceae;g__uncultured | 0.2674 | 0.5191 | 63 | 14.6 |
| d__Bacteria;p__Proteobacteria;c__Gammaproteobacteria;o__Burkholderiales;f__Comamonadaceae;g__Variovorax | 0.2459 | 0.4773 | 33.9 | 52.3 |
| d__Bacteria;p__Firmicutes;c__Bacilli;o__Bacillales;f__Bacillaceae;g__Bacillus | 0.1684 | 0.3267 | 18.2 | 34.1 |
| d__Bacteria;p__Proteobacteria;c__Gammaproteobacteria;o__Burkholderiales;f__Comamonadaceae;NA | 0.1021 | 0.1981 | 26.5 | 15.1 |

**Supplementary Material Table S5.** Differentially-abundant bacterial taxa correlated with testosterone in female northern jacanas from the MaAsLin2 analysis.

| Bacterial Taxa | Coefficient | Standard Error | p-value |
|---|---|---|---|
| d__Bacteria;p__Firmicutes;c__Bacilli;NA;NA;NA;NA | -455.67 | 8.51 | 1.64E-11 |
| d__Bacteria;p__Firmicutes;c__Bacilli;NA;NA;NA;NA | 35.16 | 0.61 | 9.01E-12 |
| d__Bacteria;p__Firmicutes;c__Bacilli;NA;NA;NA;NA | -456.44 | 8.51 | 1.62E-11 |
| d__Bacteria;p__Firmicutes;c__Bacilli;NA;NA;NA;NA | 4.43 | 0.79 | 0.00051151 |
| d__Bacteria;p__Firmicutes;c__Bacilli;NA;NA;NA;NA | 37.80 | 0.61 | 5.12E-12 |
| d__Bacteria;p__Firmicutes;c__Bacilli;NA;NA;NA;NA | 34.87 | 0.61 | 9.70E-12 |
| d__Bacteria;p__Firmicutes;c__Bacilli;o__Bacillales;f__Marinococcaceae;g__Marinococcus;NA | 33.93 | 0.61 | 1.20E-11 |
| d__Bacteria;p__Firmicutes;c__Bacilli;NA;NA;NA;NA | -440.49 | 8.51 | 2.15E-11 |
| d__Bacteria;p__Firmicutes;c__Bacilli;NA;NA;NA;NA | 33.05 | 0.61 | 1.47E-11 |
| d__Bacteria;p__Firmicutes;c__Bacilli;NA;NA;NA;NA | 34.14 | 0.61 | 1.14E-11 |
| d__Bacteria;p__Firmicutes;c__Bacilli;NA;NA;NA;NA | 36.17 | 0.61 | 7.22E-12 |
| d__Bacteria;p__Firmicutes;c__Bacilli;o__Lactobacillales;NA;NA;NA | -471.84 | 8.51 | 1.24E-11 |
| d__Bacteria;p__Firmicutes;c__Bacilli;NA;NA;NA;NA | 6.88 | 0.54 | 1.38E-06 |
| d__Bacteria;p__Firmicutes;c__Bacilli;NA;NA;NA;NA | 37.68 | 0.61 | 5.29E-12 |
| d__Bacteria;p__Firmicutes;c__Bacilli;NA;NA;NA;NA | 3.95 | 0.88 | 0.00201836 |
| d__Bacteria;p__Proteobacteria;c__Gammaproteobacteria;o__Burkholderiales;f__Comamonadaceae;g__Variovorax;s__Variovorax_paradoxus | 36.12 | 0.61 | 7.33E-12 |

| | | | |
|---|---|---|---|
| d__Bacteria;p__Firmicutes;c__Bacilli;NA;NA;NA;NA | 37.85 | 0.61 | 5.08E-12 |
| d__Bacteria;p__Firmicutes;c__Bacilli;NA;NA;NA;NA | 35.66 | 0.61 | 8.15E-12 |
| d__Bacteria;p__Firmicutes;c__Bacilli;o__Exiguobacterales;f__Exiguobacteraceae;g__Exiguobacterium;NA | 4.58 | 0.76 | 0.00032583 |
| d__Bacteria;p__Firmicutes;c__Bacilli;NA;NA;NA;NA | 4.18 | 0.83 | 0.00101813 |
| d__Bacteria;p__Firmicutes;c__Bacilli;o__Lactobacillales;NA;NA;NA | 4.70 | 0.75 | 0.00023545 |
| d__Bacteria;p__Firmicutes;c__Bacilli;NA;NA;NA;NA | 4.38 | 0.79 | 0.00056305 |
| d__Bacteria;p__Firmicutes;c__Bacilli;NA;NA;NA;NA | 5.03 | 0.70 | 9.56E-05 |
| d__Bacteria;p__Firmicutes;c__Bacilli;o__Lactobacillales;NA;NA;NA | -455.52 | 8.51 | 1.65E-11 |
| d__Bacteria;p__Firmicutes;c__Bacilli;o__Lactobacillales;f__Enterococcaceae;g__Enterococcus;s__Enterococcus_gallinarum | 35.06 | 0.61 | 9.28E-12 |
| d__Bacteria;p__Firmicutes;c__Bacilli;o__Bacillales;NA;NA;NA | -447.91 | 8.51 | 1.89E-11 |
| d__Bacteria;p__Firmicutes;c__Bacilli;o__Exiguobacterales;f__Exiguobacteraceae;g__Exiguobacterium;NA | 37.49 | 0.61 | 5.46E-12 |
| d__Bacteria;p__Firmicutes;c__Bacilli;NA;NA;NA;NA | -431.20 | 8.51 | 2.55E-11 |
| d__Bacteria;p__Firmicutes;c__Bacilli;o__Exiguobacterales;f__Exiguobacteraceae;g__Exiguobacterium;NA | -476.46 | 8.51 | 1.15E-11 |
| d__Bacteria;p__Firmicutes;c__Bacilli;NA;NA;NA;NA | 3.64 | 0.95 | 0.00491083 |
| d__Bacteria;p__Firmicutes;c__Bacilli;o__Bacillales;NA;NA;NA | -30.53 | 6.96 | 0.00232224 |
| d__Bacteria;p__Firmicutes;c__Bacilli;NA;NA;NA;NA | 35.47 | 0.61 | 8.50E-12 |
| d__Bacteria;p__Firmicutes;c__Bacilli;NA;NA;NA;NA | 32.15 | 0.61 | 1.83E-11 |
| d__Bacteria;p__Proteobacteria;c__Gammaproteobacteria;o__Burkholderiales;f__Comamonadaceae;g__uncultured;s__uncultured_bacterium | 34.47 | 0.61 | 1.06E-11 |

| | | | |
|---|---|---|---|
| d__Bacteria;p__Firmicutes;c__Bacilli;NA;NA;NA;NA | 35.66 | 0.61 | 8.15E-12 |
| d__Bacteria;p__Firmicutes;c__Bacilli;NA;NA;NA;NA | 35.60 | 0.61 | 8.22E-12 |
| d__Bacteria;p__Firmicutes;c__Bacilli;o__Exiguobacterales;f__Exiguobacteraceae;g__Exiguobacterium;NA | -477.48 | 8.51 | 1.13E-11 |
| d__Bacteria;p__Firmicutes;c__Bacilli;NA;NA;NA;NA | -433.04 | 8.51 | 2.47E-11 |
| d__Bacteria;p__Firmicutes;c__Bacilli;o__Exiguobacterales;f__Exiguobacteraceae;g__Exiguobacterium;NA | 35.79 | 0.61 | 7.92E-12 |
| d__Bacteria;p__Firmicutes;c__Bacilli;o__Lactobacillales;f__Enterococcaceae;g__Enterococcus;s__Enterococcus_gallinarum | -439.01 | 8.51 | 2.21E-11 |
